# Metagenomic Insights into the Fecal Microbiome, Resistome, and Mobilome of Beef Cattle in Cow–Calf Operations in Florida, United States

**DOI:** 10.64898/2026.11.04.764209

**Authors:** Ahmad Ali, Lekshmi K Edison, João H. J. Bittar, Thomas Denagamage, Alice C. Y. Lee, Kwang C. Jeong, Jorge A. Hernandez, Subhashinie Kariyawasam

## Abstract

Food-producing animals are a focus of One Health approaches to antimicrobial resistance (AMR) as reservoirs of antimicrobial-resistant bacteria (ARB) and antimicrobial resistance genes (ARGs). We hypothesized that microbial composition, ARGs profiles, and mobile genetic element (MGE) profiles would differ by operation size and animal age group. Fecal samples from 162 cows and calves were combined into 54 composite samples, each comprising three animals of the same age group. Across operation sizes, *Bacillota* (51.4–64.9%), *Bacteroidota* (12.0–17.3%), *Pseudomonadota* (10.3–16.5%), and *Actinomycetota* (3.3–10.9%) were predominant phyla. Alpha diversity did not differ among operation sizes; however, cows exhibited greater observed richness, Shannon diversity, and Simpson diversity than calves (all *P* < 0.001). Microbial composition did not differ significantly among operation sizes (PERMANOVA, R² = 0.065, *P* = 0.609) but differed significantly between cows and calves (R² = 0.200, *P* < 0.001). Seventy-five ARGs representing 14 resistance classes were identified. Normalized ARG hit rates were higher in small operations (0.444) than in medium (0.278) and large operations (0.222; *P* = 0.088), whereas calves exhibited a higher estimated marginal mean normalized ARG hit rate than cows (0.383 vs. 0.247 hits/million reads; *P* < 0.001). Of 1,054 curated ARG hits, 23.9% occurred on predicted plasmid contigs, 0.9% on predicted viral contigs, and 7.1% on contigs containing insertion sequences. Overall findings demonstrate differences in fecal microbiome diversity and ARG abundance between cows and calves and indicate that predicted plasmids are associated with a notable proportion of ARG hits in Florida beef cow–calf operations.

**IMPORTANCE:** Beef cow–calf operations are an important part of beef cattle production, yet relatively little is known about the microorganisms and antimicrobial resistance genes present in cattle feces in these production systems. This study examined fecal microbial communities, antimicrobial resistance genes, and genetic elements associated with their potential movement in beef cows and calves from nine Florida beef cow–calf operations. We found that cows and calves differed in their microbial communities, with cows showing greater microbial diversity and calves exhibiting higher abundance of antimicrobial resistance genes. In contrast, operation size was not significantly associated with these outcomes. Nearly one-quarter of the detected resistance gene hits occurred on DNA fragments predicted to originate from plasmids, which can facilitate gene transfer between bacteria. These findings improve our understanding of antimicrobial resistance in beef cow–calf production and provide information relevant to monitoring resistance in food-producing animals.

## 1. INTRODUCTION

Antimicrobial resistance (AMR) poses a major global public health concern by compromising the effective treatment of bacterial infection (1, 2). In 2021, bacterial AMR was associated with more than one million deaths worldwide, and its global burden is projected to increase considerably over the coming years if effective mitigation strategies are not implemented (3). Food-producing animals are a key focus of One Health approaches to AMR because they can serve as reservoirs of antimicrobial-resistant bacteria (ARB) and antimicrobial resistance genes (ARGs) (4, 5). These bacteria and resistant genes may disseminate among humans, animals, and the environment through direct and indirect pathways via the food chain, and contamination of soil, and water by animal feces. This interconnectedness highlights the importance of investigating AMR in livestock production systems through a One Health approach (5, 6).

The gastrointestinal (GI) microbiome plays an essential role in host health across mammalian species by contributing to metabolism, immune development, and colonization resistance against enteric pathogens (7). The gut microbiome also serves as a major reservoir of ARGs, collectively referred to as the resistome. The dissemination and persistence of these resistance genes are facilitated by mobile genetic elements (MGEs), including plasmids, bacteriophages, and insertion sequences (IS) as well as associated gene-capture systems such as integrons (In), which collectively contribute to the acquisition and spread of resistance among bacterial populations (8). Therefore, integrated characterization of the microbiome, resistome, and mobilome provides a more comprehensive understanding of AMR ecology than approaches that focus exclusively on cultured bacteria or on individual resistance determinants.

The United States is among the world’s leading beef-producing countries. Cow–calf operations underpin the U.S. beef production system by supplying weaned calves for subsequent stocker and feedlot operations (9, 10). Cow-calf operations are typically managed under extensive, pasture-based production systems, where prophylactic and metaphylactic antimicrobial use, as well as disease requiring antimicrobial treatment, may occur less frequently than in more intensive production systems (11, 12). Differences in management practices, stocking densities, and antimicrobial use patterns may contribute to variations in the fecal microbiome and AMR profiles between beef cow–calf operations and feedlots. Florida is predominantly a cow-calf state and ranks among the top ten U.S. states in beef cow inventory (13). Its subtropical climate, year-round grazing, and predominantly pasture-based management practices provide distinct production environments that may uniquely shape the fecal microbiome, resistome, and mobilome of beef cattle.

Previous studies of AMR in beef cow–calf operations have primarily relied on culture-based methods and targeted phenotypic or genotypic analyses of selected bacterial species. Agga et al. (14) investigated antimicrobial-resistant fecal bacteria in beef cows using culture-based methods and targeted molecular detection of resistance genes. Similarly, Morris et al. (15) examined antimicrobial resistance in fecal *Escherichia coli* and *Enterococcus* spp. from beef cow–calf operations in northern California using culture-based isolation and antimicrobial susceptibility testing. Furthermore, a systematic review by Ali et al. (16) highlighted that previous studies of antimicrobial-resistant fecal bacteria in beef cow–calf operations largely relied on culture-based phenotypic susceptibility testing of selected bacterial species, particularly *Escherichia coli* and *Enterococcus* spp., with some studies incorporating targeted molecular methods. Although previous studies have provided valuable insights into AMR in selected bacterial populations, integrated characterization of the fecal microbiome, resistome, and mobilome in beef cow–calf operations remains limited, particularly across operation sizes and between cows and calves. Differences in antimicrobial use, management practices, and environmental exposure across operations, as well as diet, gastrointestinal development, and immune maturation between cows and calves, may influence microbial community composition and ARG distribution. To address these knowledge gaps, this study aimed to characterize the fecal microbiome, resistome, and mobilome in beef cow–calf operations in Florida using shotgun metagenomic sequencing. Specifically, we evaluated differences in microbial community composition and ARG abundance and distribution across operation sizes and animal groups (cows and calves) and examined associations between ARGs and MGEs. We hypothesized that fecal microbial community composition, ARG abundance, and distribution would differ by operation size and animal group.

## 2. MATERIALS AND METHODS

### 2.1 Study Cow-Calf Operations

Nine beef cow-calf operations located in five counties in Florida (Hardee, Marion, Okeechobee, Pasco, and Sumter) were included in the study. Based on the herd size, the operations were categorized as small (20–49 cows), medium (50–199 cows), and large (≥200 cows) (17). Three operations were selected from each herd size category (n=3 per category). Animal sampling procedures were approved by the University of Florida Institutional Animal Care and Use Committee (IACUC #202300000165).

### 2.2 Sample Selection and Pooling Strategy

A total of 162 fecal samples were selected from 743 samples collected in a previous study evaluating AMR bacteria in beef cattle from cow-calf operations (18). From each of the nine beef cow calf operations, fecal samples from nine cows and nine calves were randomly selected using an online random number generator (Research Randomizer) to minimize selection bias. Within each operation, nine cow samples were randomly assigned into three pools (three cows per pool). The nine calf samples were pooled separately using the same approach (three calves per pool).

This resulted in three cow pools and three calf pools per operation, yielding six pooled samples per operation and a total of 54 pooled fecal samples (9 operations × 6 pools) for DNA extraction and subsequent shotgun metagenomic sequencing (Figure 1).

**Figure 1.**
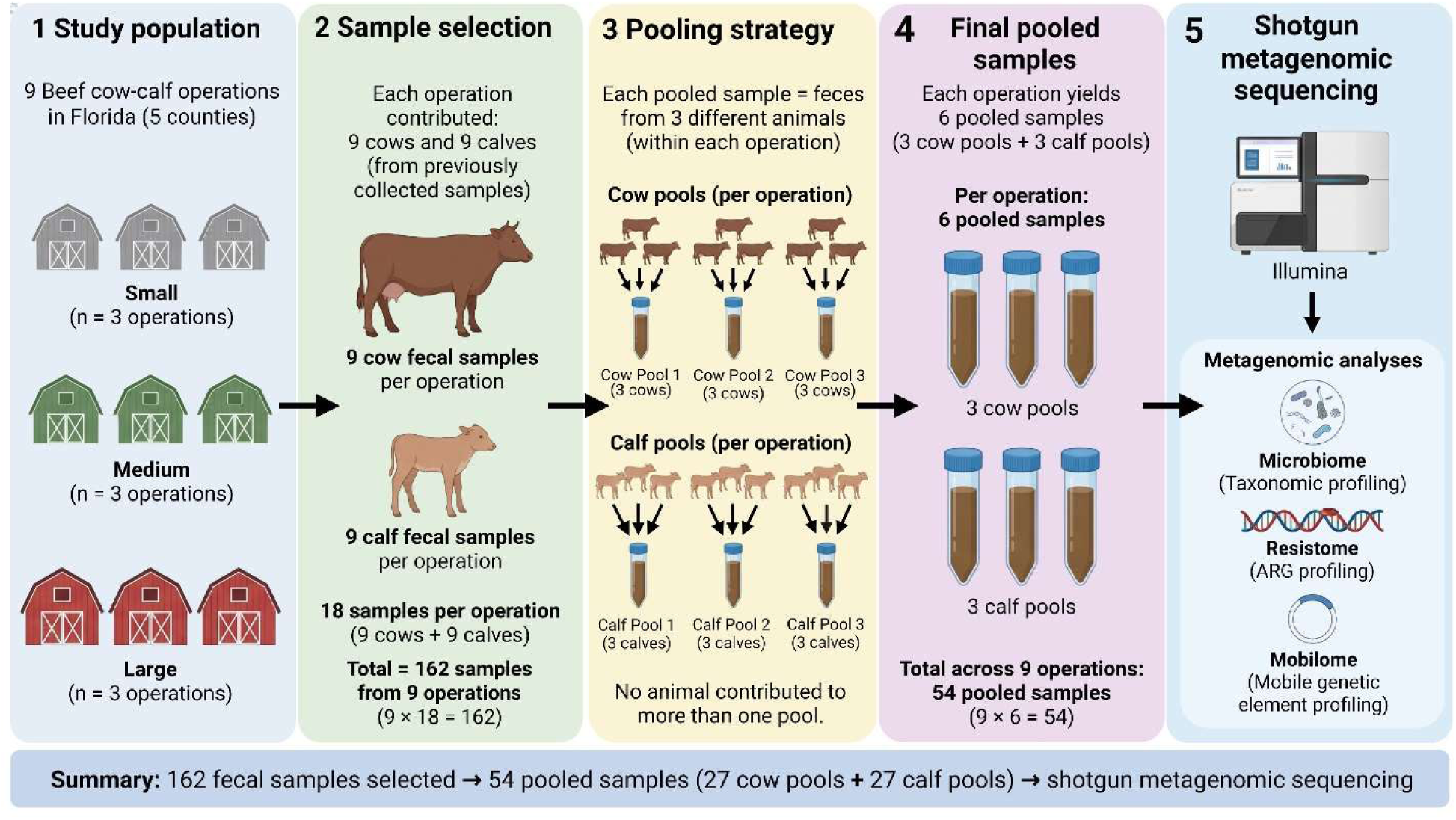
Overview of the study design and shotgun metagenomic workflow.

### 2.3 DNA extraction and sequencing

DNA was extracted using the QIAamp PowerFecal Pro DNA Kit (QIAGEN, Germantown, MD, USA) according to the manufacturer’s instructions. DNA purity was initially assessed using a NanoDrop spectrophotometer (Thermo Fisher Scientific, Waltham, MA, USA), and DNA concentration was quantified using a Qubit fluorometer (Thermo Fisher Scientific). Extracted DNA was stored at −80°C until library preparation. Sequencing libraries were prepared using the Illumina DNA Prep Kit (Illumina, San Diego, CA, USA), and sample indexing was performed using the Illumina DNA/RNA UD Indexes Set A, Tagmentation (Illumina, San Diego, CA, USA), according to the manufacturer’s instructions. Prepared libraries were submitted to the University of Florida Interdisciplinary Center for Biotechnology Research (UF-ICBR) for shotgun metagenomic sequencing.

### 2.4 Bioinformatics Analysis

Bioinformatic analyses of shotgun metagenomic datasets were performed using the University of Florida HiPerGator high-performance computing cluster. The raw paired-end sequencing reads were quality assessed using FastQC v0.12.1. Adapter sequences and low-quality bases were removed using Trimmomatic v0.39. Host-derived reads were removed by aligning quality-filtered reads against the *Bos taurus* reference genome (ARS-UCD1.2; Ensembl release 110) using Bowtie2 v2.5.4. Read pairs in which both mates remained unmapped to host reference genome were retained using SAMtools v1.20 for all downstream analyses.

For microbiome characterization, taxonomic classification was performed with Kraken2 v2.17.1 against the K2 Standard database (19), followed by abundance re-estimation using Bracken v2.9. Taxonomic profiles were generated at the phylum and genus levels, and the relative abundance of each taxon was calculated as the percentage of reads assigned to that taxon within each sample. Alpha diversity was evaluated at the genus level using observed richness, Shannon diversity, and Simpson diversity indices. Beta diversity was assessed using Bray–Curtis dissimilarities and visualized using non-metric multidimensional scaling (NMDS).

For resistome characterization, host-filtered reads from each sample were assembled *de novo* using MEGAHIT (20). Contigs ≥1,000 bp were retained for downstream analyses. ARGs were identified using NCBI AMRFinderPlus v4.2.7 and classified according to their respective ARG classes (21). To account for differences in sequencing depth among samples, the number of ARG hits identified in each sample was normalized to the total number of sequencing reads and expressed as ARG hits per million reads [(number of ARG hits/total number of reads) × 10⁶].

To characterize the genomic context and potential mobility of ARGs, assembled contigs were analyzed using geNomad v1.5.1 to identify plasmids and viruses (22), IntegronFinder v2.0.6 to identify integron-related structures (23), and ISEScan v1.7.3 to identify insertion sequences (ISs) (24). Predictions from these tools were linked to the curated ARG dataset using shared sample and contig identifiers. ARGs located on contigs predicted as plasmids or virus were classified as plasmid-or virus-associated, respectively. Integron-related structures were categorized as complete integrons, In0 elements, or clusters of *attC* (attachment site of the cassette) lacking an integron-integrase gene (CALINs). IS-containing contigs were identified using the default ISEScan settings, retaining both complete and partial IS predictions. For ARGs located on IS-containing contigs, genomic coordinates were compared with those of the predicted ISs to assess whether their sequences were overlapped. Non-overlapping ARGs were further categorized based on their distance from the nearest predicted IS (≤100 bp, 101–1,000 bp, or >1,000 bp). When multiple ISs were identified on the same contig, overlapping ISs were prioritized; otherwise, the nearest IS was retained for ARG-level analysis.

### 2.5 Statistical Analysis

All statistical analyses and data visualization were performed in R version 4.3 using RStudio. Statistical analyses were conducted to evaluate differences in fecal microbiome and resistome profiles according to operation size and animal group. For microbiome analyses, alpha-diversity indices including, observed richness, Shannon diversity, and Simpson diversity, were analyzed using linear mixed-effects models. Operation size (small, medium, and large) and animal group (cow and calf) were included as fixed effects, with farm included as a random intercept to account for clustering of samples within operations. Beta diversity was quantified using Bray– Curtis dissimilarity and visualized by non-metric multidimensional scaling (NMDS). Differences in microbial community composition were evaluated using permutational multivariate analysis of variance (PERMANOVA) implemented in the vegan package, with operation size and animal group analyzed separately. The proportion of variation explained (R²) and corresponding *P* values were reported. For resistome analyses, normalized ARG hit rates, expressed as ARG hits per million reads, were analyzed using linear mixed-effects models. Operation size and animal group were included as fixed effects, and farm was included as a random intercept. Statistical significance was defined as a *P* value < 0.05. Descriptive statistics were used to summarize the genomic associations of ARGs with predicted mobilome features, including plasmid-associated, viral-associated, and insertion sequence-containing ARGs, both overall and according to operation size and animal group.

## 3. RESULTS

### 3.1 Shotgun Metagenomic Sequencing

Shotgun metagenomic sequencing was performed on all 54 pooled fecal samples, collected from nine beef cow–calf operations. Sequencing generated a total of 3.54 billion paired-end reads, with a mean sequencing depth of approximately 65.5 million reads per sample (ranging from 15.3 to 105.0 million reads). Raw sequencing read counts for individual samples are provided in Supplementary Table S1. Following quality filtering and removal of host-derived reads, the remaining reads were used for microbiome profiling and *de novo* assembly. The resulting assembled contigs were subsequently used for resistome, and mobilome analyses.

### 3.2 Phylum-level Microbial Composition

The fecal microbiome was predominantly composed of four bacterial phyla: *Bacillota*, *Bacteroidota*, *Pseudomonadota*, and *Actinomycetota* (Figure 2A-C). Across operation sizes, *Bacillota* was the most abundant phylum, representing 51.4%, 64.2%, and 64.9% of the microbial community in small, medium, and large operations, respectively (Figure 2B). The relative abundance of *Bacteroidota* decreased with increasing operation size, from 17.3% in small operations to 15.8% in medium operations and 12.0% in large operations. Similarly, the relative abundance of *Pseudomonadota* and *Actinomycetota* decreased with increasing operation size from accounted for 16.5% and 10.9% in small operations, from 10.6%, and 6.1% in medium operations, and 10.3% and 3.3% in large operations, respectively. Among the less abundant phyla, *Verrucomicrobiota* exhibited a higher relative abundance in large operations (3.8%) than in medium (0.2%) or small (0.1%) operations. *Campylobacterota* accounted for 1.7% of the microbial community in large operations compared with less than 1% in other operation sizes. At the animal age group level, *Bacillota* was the predominant phylum in both calves and cows, accounting for 59.1% and 61.3% of the microbial community, respectively (Figure 2C). *Bacteroidota* exhibited a higher relative abundance in calves (17.2%) than in cows (12.9%), whereas *Actinomycota* was more abundant in cows (9.3%) than calves (4.2%). *Pseudomonadota* showed comparable relative abundances in calves (12.5%) and cows (12.4%). Among the less abundant phyla, *Verrucomicrobiota* exhibited a higher relative abundance in calves (2.6%) than in cows (0.1%). Variation in the relative abundance of individual phyla was also observed across operations and animal groups (Figure 2A). Complete phylum-level relative abundance data for all 54 pooled samples are provided in Supplementary Table S2.

**Figure 2.**
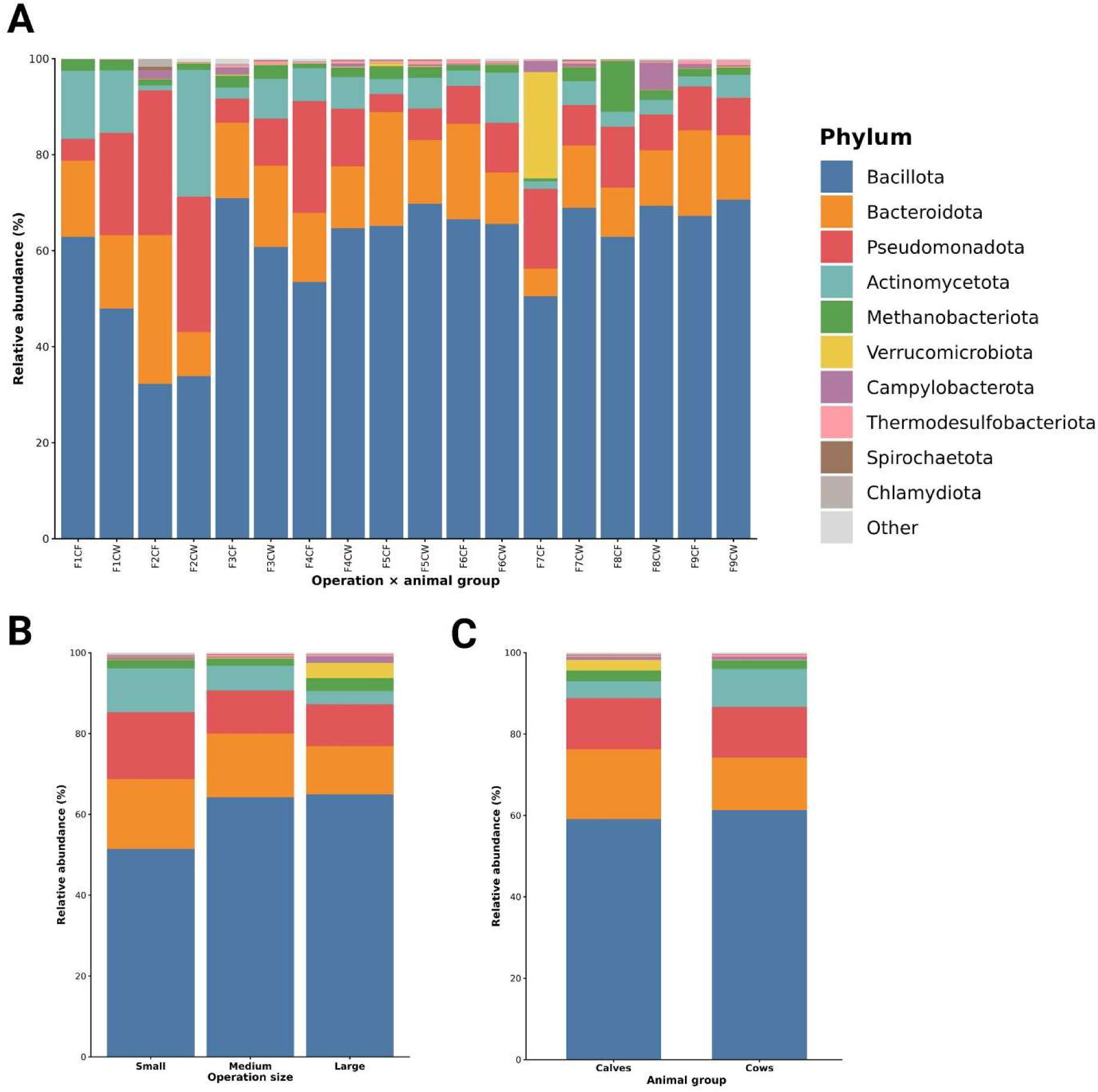
Phylum-level taxonomic composition of the fecal microbiome in beef cow–calf operations. **(A)** Relative abundance of bacterial phyla in pooled fecal samples grouped by operation size; CF denotes calves, and CW denotes cows. **(B)** Relative abundance of bacterial phyla according to operation size. **(C)** Relative abundance of bacterial phyla according to animal type. Taxa not displayed individually were grouped as "Other."

### 3.3 Genus-level Microbial Composition

Relative abundance profiles showed greater taxonomic variability at the genus level than at the phylum level (Figure 3A–C). Although numerous bacterial genera were detected, a relatively small number of dominant genera accounted for a substantial proportion of the microbial community, whereas the remaining low-abundance genera were grouped as “Other.” Across operation sizes, *Lactobacillus, Escherichia, Blautia, Bacteroides, Romboutsia,* and *Streptococcus* were among the predominant genera (Figure 3B). *Lactobacillus* exhibited a higher relative abundance in medium (15.1%) and large (14.7%) operations than in small operations (8.3%). *Escherichia* accounted for 7.4%, 7.1%, and 8.3% of the microbial community in small, medium, and large operations, respectively, whereas *Blautia* showed comparable relative abundances across operation sizes (6.1%, 5.9%, and 6.3%, respectively). *Streptococcus* exhibited a higher relative abundance in large operations (7.7%) than in small (3.4%) and medium (3.3%) operations. Among the other genera, *Akkermansia* accounted for 3.8% of the microbial community in large operations compared with less than 0.2% in small and medium operations. Similarly, *Campylobacter* represented 1.7% in large operations, compared with less than 1% in the other operation sizes. Genus-level microbial composition also differed between cows and calves (Figure 3C). *Lactobacillus* was the predominant genus in calves, with a higher relative abundance in calves than in cows (20.2% vs. 5.2%). Similarly, *Escherichia* (11.3% vs.3.9%), *Limosilactobacillus* (5.4% vs. 2.1%) and *Akkermansia* (2.6% vs. 0.1%) exhibited higher relative abundances in calves than in cows. In contrast, *Romboutsia* (8.8% vs. 2.4%) and *Bacillus* (5.2% vs. 0.4%) had higher relative abundances in cows than in calves. *Blautia* (6.0% in cows vs. 6.2% in calves) and *Bacteroides* (5.7% in both groups) showed comparable relative abundances between the cows and calves. Complete genus-level relative abundance data for all 54 pooled samples are provided in Supplementary Table S3.

**Figure 3.**
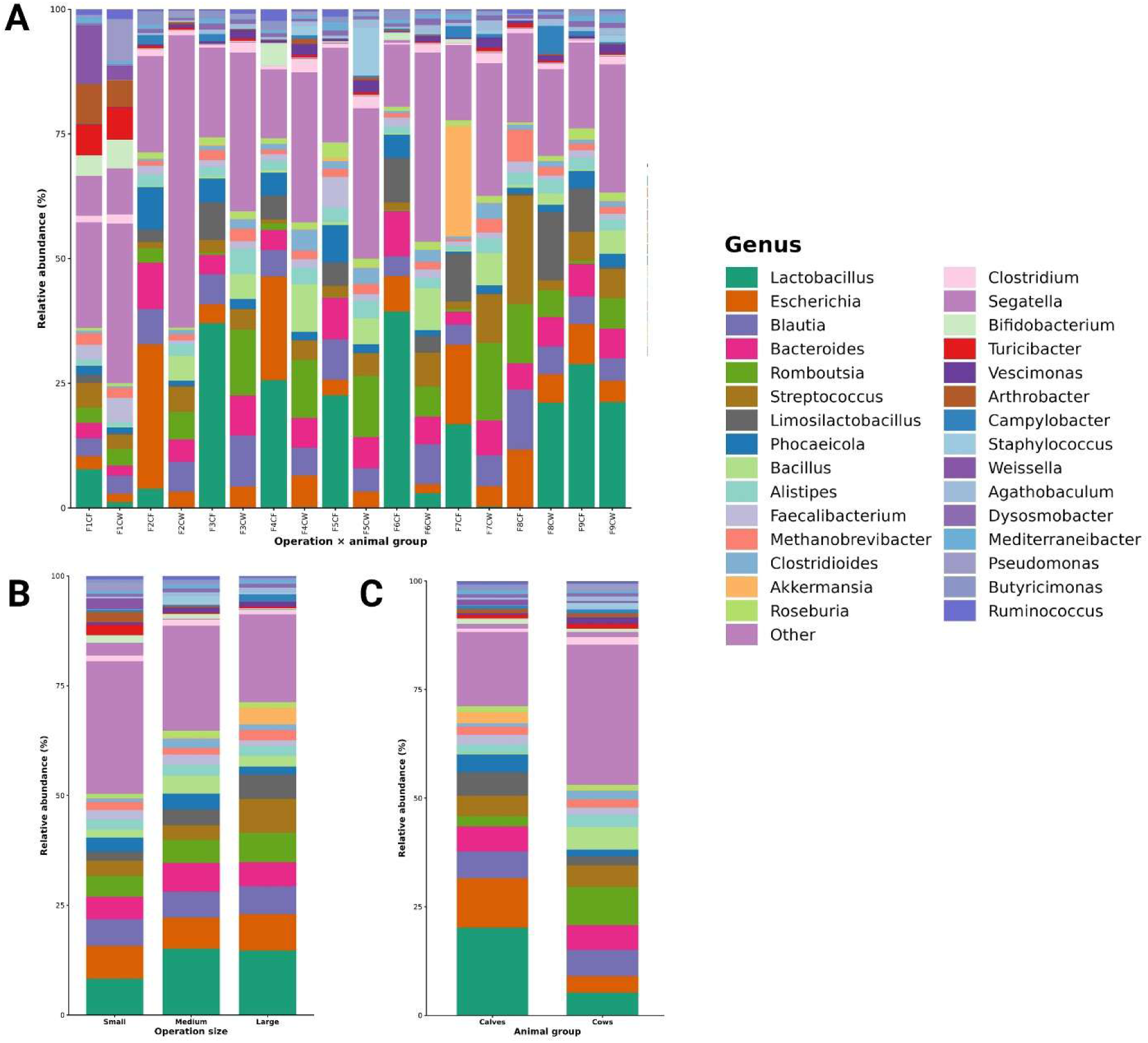
Genus-level taxonomic composition of the fecal microbiome in beef cow–calf operations. **(A)** Relative abundance of bacterial genera in pooled fecal samples grouped by operation size; CF denotes calves, and CW denotes cows. **(B)** Relative abundance of bacterial genera according to operation size. **(C)** Relative abundance of bacterial genera according to animal type. Taxa not displayed individually were grouped as "Other."

### 3.4 Alpha Diversity

Alpha diversity was evaluated at the genus level using observed richness, Shannon diversity, and Simpson diversity indices. No statistically significant differences were observed among operation-size categories for observed richness (*P* = 0.343) and Simpson diversity (*P* = 0.157), while Shannon diversity showed some evidence of differences (*P* = 0.069). (Figure 4A–C). In contrast, all three alpha-diversity indices differed significantly between cows and calves (Figure 5A–C). Cows exhibited greater observed genus richness than calves (estimated marginal means: 1,280 vs. 1,143 genera; *P* < 0.001), as along with higher Shannon diversity (3.72 vs. 2.90; *P* < 0.001) and Simpson diversity (0.929 vs. 0.860; *P* < 0.001). These findings indicate greater genus-level microbial richness and diversity in cows than in calves, with no statistically significant differences detected among operation sizes.

**Figure 4.**
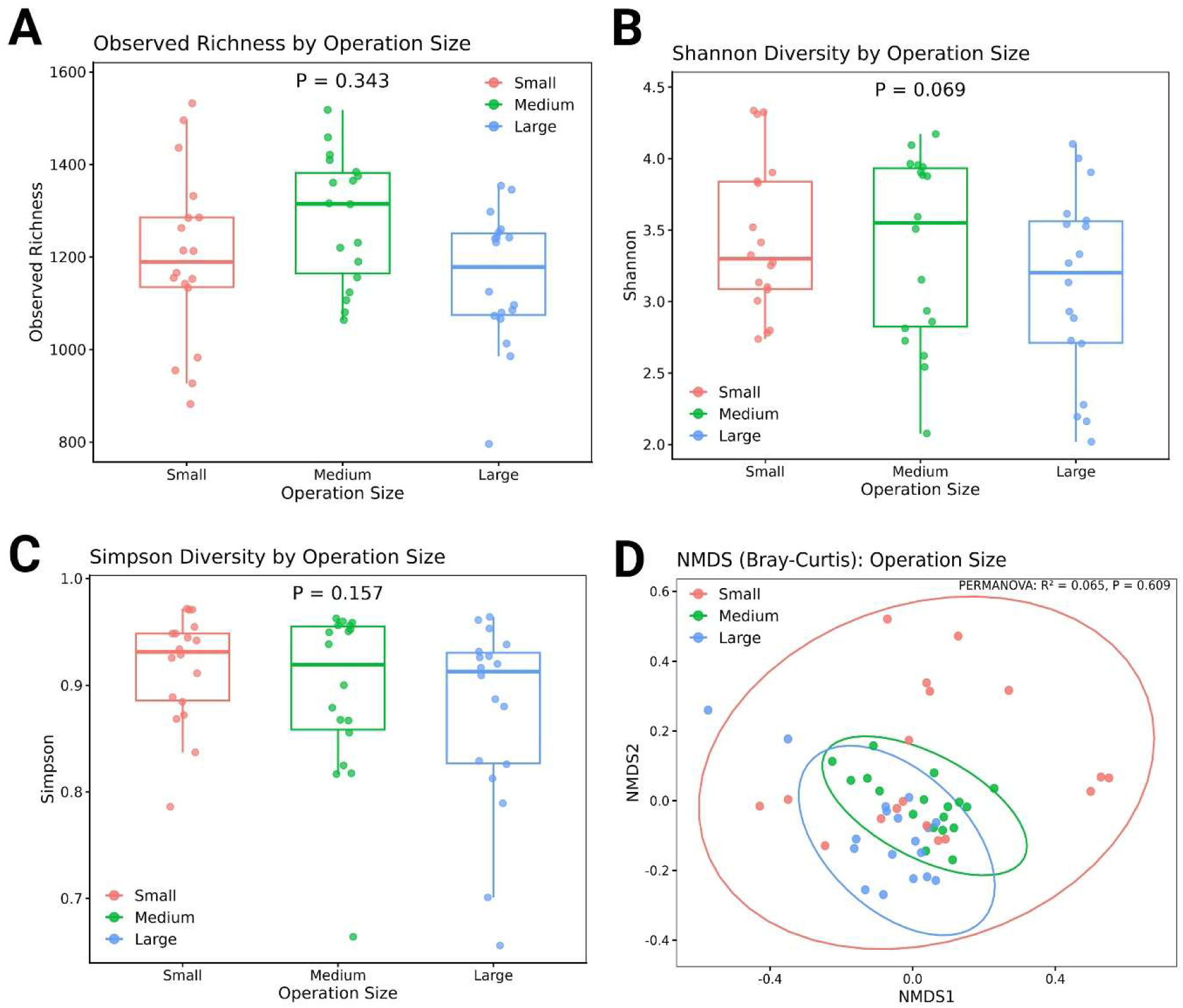
Alpha and beta diversity of the fecal microbiome by operation size. **(A)** Observed richness. **(B)** Shannon diversity. **(C)** Simpson diversity. **(D)** Non-metric multidimensional scaling (NMDS) based on Bray–Curtis dissimilarity.

**Figure 5.**
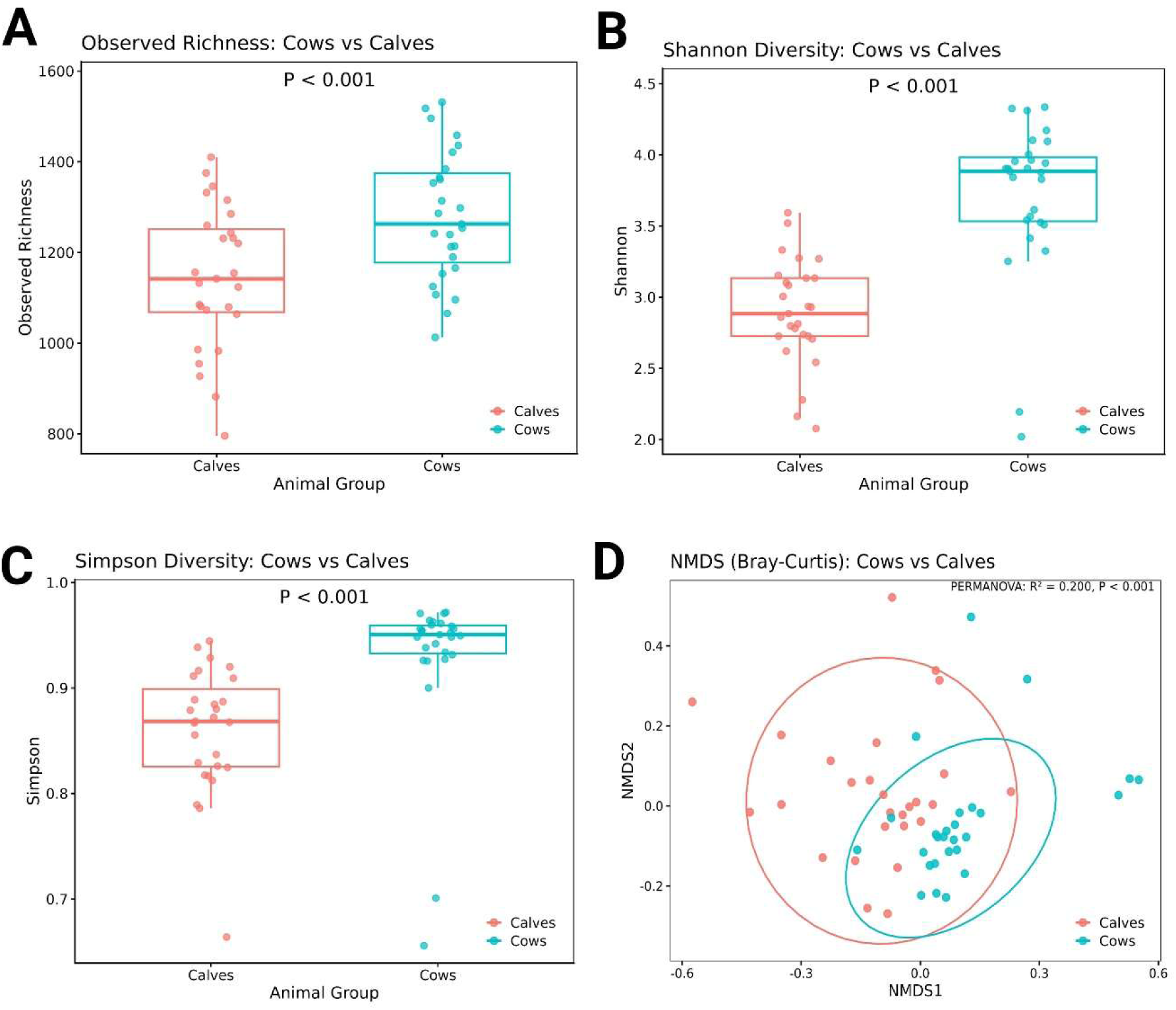
Alpha and beta diversity of the fecal microbiome according to animal group. **(A)** Observed richness. **(B)** Shannon diversity. **(C)** Simpson diversity. **(D)** Non-metric multidimensional scaling (NMDS) based on Bray–Curtis dissimilarity.

### 3.5 Beta Diversity

Beta diversity was assessed using Bray–Curtis dissimilarity and visualized by non-metric multidimensional scaling (NMDS). Microbial community composition did not differ significantly among operation size categories (PERMANOVA, *R*² = 0.065, *P* = 0.609; Figure 4D). However, a significant difference was observed between cows and calves (PERMANOVA, *R*² = 0.200, *P* < 0.001; Figure 5D). Animal group explained a greater proportion of variation in microbial community composition (20.0%) than operation size (6.5%). Overall, these findings suggest that animal groups were significantly associated with differences in fecal microbial community composition, whereas operation size was not.

### 3.6 Resistome Analysis

ARGs were detected in all 54 pooled metagenomic samples. A total of 75 unique ARGs representing 14 AMR classes were identified, with genes conferring resistance to tetracycline, aminoglycosides, β-lactams, macrolides, and lincosamides comprising the predominant components of the resistome (Figure 6). Overall, 1,054 ARG hits were identified across all samples. Detailed ARG-level results for each sample are provided in Supplementary Table S4.

**Figure 6.**
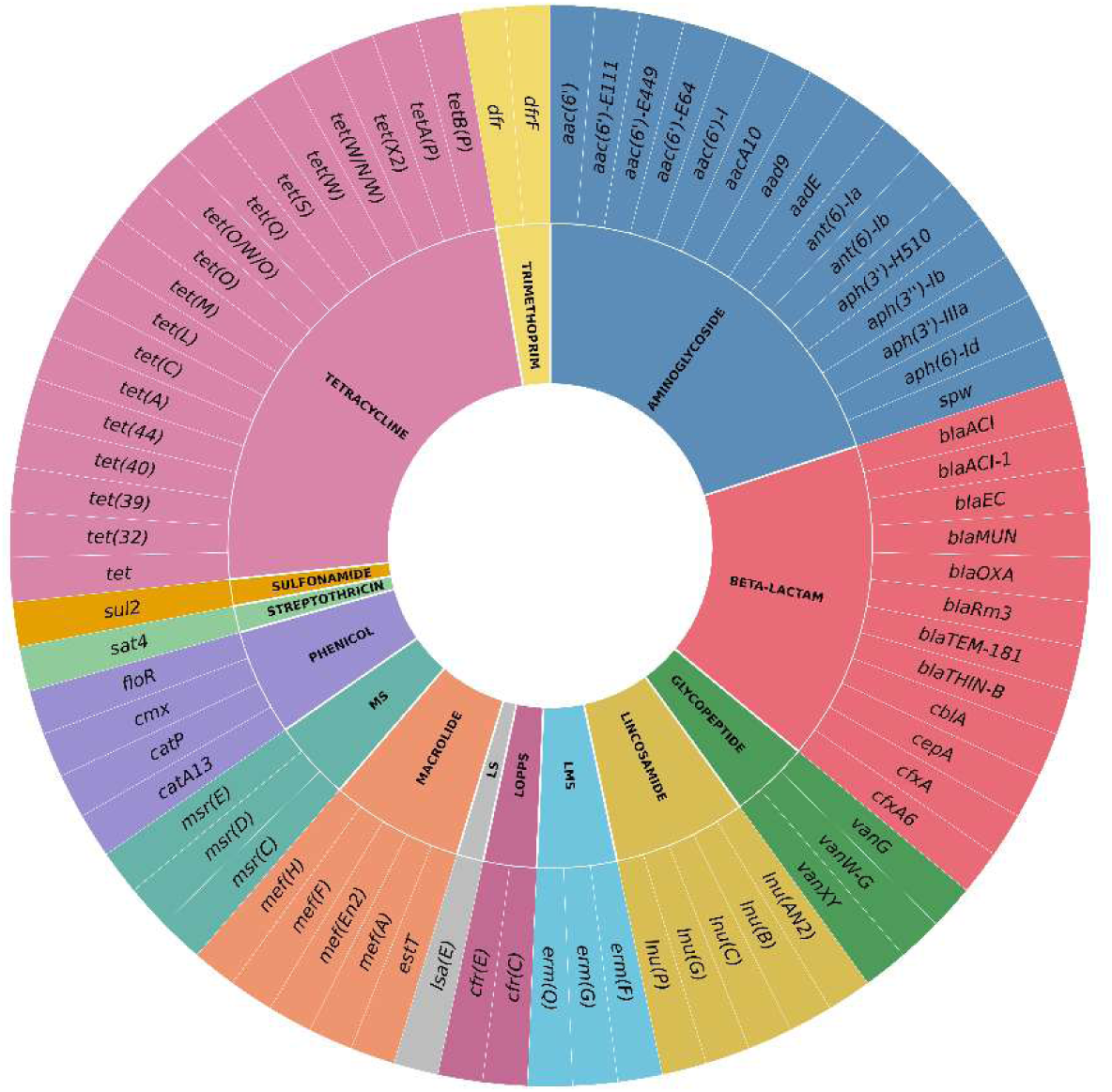
Overall diversity of AMR genes identified in the study population. The sunburst plot illustrates the distribution of detected ARGs according to antimicrobial class. The inner ring represents AMR classes, while the outer ring represents individual resistance genes. (LMS = Lincosamide/Macrolide/Streptogramin; LOPPS = Lincosamide/Oxazolidinone/Phenicol/Pleuromutilin/Streptogramin; LS = Lincosamide/Streptogramin; MS = Macrolide/Streptogramin).

### 3.7 Distribution of ARGs According to Operation Size and Animal Age Group

The distribution of individual ARGs varied across operation sizes and between animal age groups (Figure 7). Of the 75 unique ARGs identified, 28 genes were shared among small, medium, and large operations. An additional 5, 8, and 2 ARGs were shared exclusively between small and medium, small and large, and medium and large operations, respectively. In addition, 21, 6, and 5 ARGs were detected exclusively in small, medium, and large operations, respectively. Between animal groups, 43 ARGs were shared by cows and calves, whereas 11 ARGs were detected exclusively in cows and 21 exclusively in calves. Tetracycline resistance genes showed the broadest distribution across both operation sizes and animal age groups.

**Figure 7.**
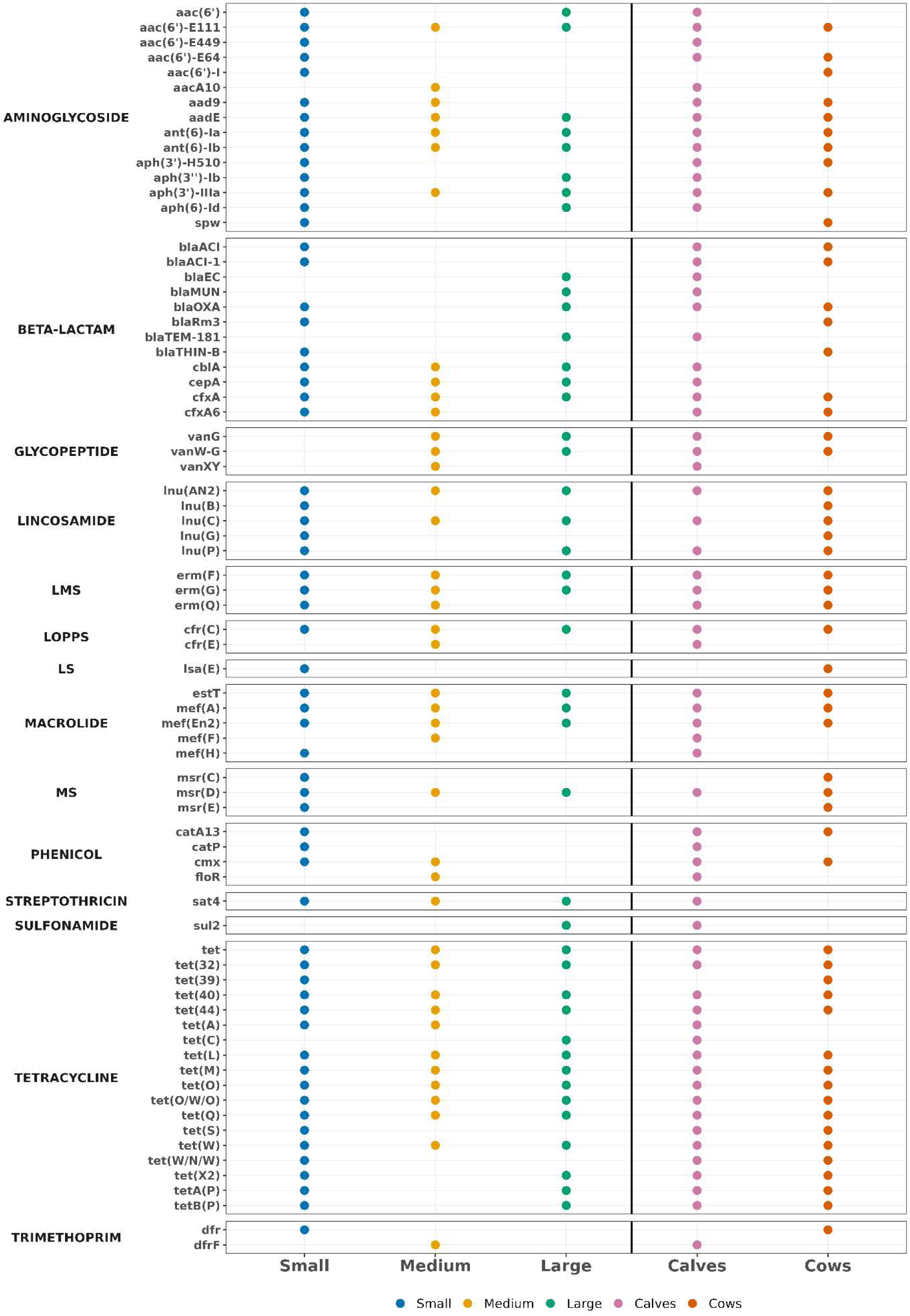
Distribution of ARGs according to operation size and animal group. The dot matrix shows the presence of individual ARGs across small, medium, and large beef cow–calf operations, as well as in calves and cows. A dot indicates the detection of at least one gene within the corresponding group. (LMS = Lincosamide/Macrolide/Streptogramin; LOPPS = Lincosamide/Oxazolidinone/Phenicol/Pleuromutilin/Streptogramin; LS = Lincosamide/Streptogramin; MS = Macrolide/Streptogramin).

### 3.8 Overall Normalized ARG Abundance

The overall mean normalized ARG hit rate was 0.315 ARG hits per million reads. The estimated marginal mean normalized ARG hit rate was highest in small operations (0.444 ARG hits/million reads), followed by medium (0.278 ARG hits/million reads) and large (0.222 ARG hits/million reads) operations. However, the linear mixed-effects model, with farm included as a random intercept, showed no statistically significant association between operation size and normalized ARG hit rate (*P* = 0.088). In contrast, normalized ARG hit rates differed significantly between animal age groups (P < 0.001), with calves exhibiting a higher estimated marginal mean than cows (0.383 vs. 0.247 ARG hits/million reads) (Table 1).

**Table 1.** Estimated marginal mean (EMM) normalized antimicrobial resistance gene (ARG) hit rates by operation size and animal age group in cow–calf operations.

| **Factor** | **Group** | **n** | **EMM hits/million reads** | ***P*-value** |
|---|---|---|---|---|
| Operation size | Small | 18 | 0.444 | 0.088 |
|  | Medium | 18 | 0.278 |  |
|  | Large | 18 | 0.222 |  |
| Animal groups | Calf | 27 | 0.383 | <0.001 |
|  | Cow | 27 | 0.247 |  |

Note: n represents the number of pooled fecal samples. Each operation-size category included three operations and 18 pooled samples. Each animal group included 27 pooled samples. *P* values were obtained from linear mixed-effects models with farm included as a random intercept. Each *P* value represents the comparison for the corresponding factor, not an individual group.

### 3.9 Mobilome

Among the 1,054 curated ARG hits identified across the 54 pooled samples subjected to metagenomic analysis, 252 (23.9%) were located on contigs predicted as plasmids by geNomad, whereas 9 (0.9%) were located on predicted viral contigs (Table 2). Plasmid-associated ARG hits were detected across all operation sizes, representing 22.0%, 26.4%, and 24.5% of ARG hits in small, medium, and large operations, respectively. Similarly, plasmid-associated ARG hits accounted for 23.5% and 24.2% of the total ARG hits in cows and calf samples, respectively. In contrast, virus-associated ARG hits were relatively uncommon across all groups, accounting for 0.3–1.4% of ARG hits. Plasmid-associated ARGs represented multiple AMR classes, including tetracyclines, aminoglycosides, β-lactams, macrolides, lincosamides, phenicols, and sulfonamides. Detailed ARG-level associations with predicted plasmid and viral contigs are provided in Supplementary Table S4. IntegronFinder identified 2,414 clusters of CALINs, five complete integrons, and three In0 elements across the 54 pooled metagenomic samples. However, none of the 1,054 curated ARG hits were located on contigs containing any of these integron-related structures, indicating that no ARG-co-localization was detected in the assembled contigs.

**Table 2.** Distribution of ARG-associated mobilome features according to operation size and animal group.

| **ARG-associated<br>mobilome feature** | **Overall<br>n (%)** | **Operation type** |  |  | **Animal age group** |  |
|---|---|---|---|---|---|---|
|  |  | **Small<br>n (%)** | **Medium<br>n (%)** | **Large<br>n (%)** | **Cow<br>n (%)** | **Calf<br>n (%)** |
| **Plasmid- ARGs** | 252 (23.9) | 103 (22) | 75 (26.4) | 74 (24.5) | 99 (23.5) | 153 (24.2) |
| **Viral- ARGs** | 9 (0.85) | 4 (0.9) | 4 (1.4) | 1 (0.3) | 2 (0.5) | 7 (1.1) |
| **IS-ARGs** | 75 (7.1) | 33 (7.1) | 21 (7.4) | 21 (7.0) | 22 (5.2) | 53 (8.4) |
| **Integron-ARGs** | 0 (0.0) | 0 (0.0) | 0 (0.0) | 0 (0.0) | 0 (0.0) | 0 (0.0) |

Notes: Values are presented as the number of ARG hits (n) and the corresponding percentage (%). Percentages were calculated using the total number of curated ARG hits within each category (overall, operation size, or animal age groups) as the denominator. A total of 1,054 curated ARG hits were identified across 54 pooled fecal samples.

A total of 75 ARG hits (7.1% of the 1,054 curated ARG hits) were located on contigs containing at least one predicted IS identified by ISEScan. The proportion of ARG hits on IS-containing contig was comparable across operation sizes, ranging from 7.0% to 7.4%, but was descriptively higher in calves (8.4%) than in cows (5.2%) (Table 2). Of these 75 ARG hits, 10 (13.3%) directly overlapped a predicted IS, whereas 13 (17.3%), 23 (30.7%), and 29 (38.7%) were located within 100 bp, 101–1,000 bp, and more than 1,000 bp from the nearest predicted IS, respectively.

Overall, direct ARG–IS overlap was observed for 10 of the 1,054 curated ARG hits (0.95%). A summary of ARG associations with predicted MGEs is presented in Table 2, and detailed ARG-level results are provided in Supplementary Table S4.

## 4. DISCUSSION

The present study provides new insights into the fecal microbiome, resistome, and mobilome of beef cow–calf operations in Florida. Cows exhibited greater genus-level microbial richness and diversity, whereas calves had higher normalized antimicrobial resistance gene (ARG) hit rates. Although normalized ARG hit rates varied numerically across operation sizes, the differences were not statistically significant. By examining these microbial characteristics across nine operations representing three size categories and comparing cows and calves using shotgun metagenomic sequencing, this study provides a broader understanding of microbial community composition and ARG distribution in cow–calf production systems.

### 4.1 Microbiome

The fecal microbiome of cows and calves was predominantly composed of four phyla: *Bacillota, Bacteroidota, Pseudomonadota*, and *Actinomycetota*, with *Bacillota* being the most abundant across operation sizes and animal age groups. These findings are consistent with previous studies of cattle fecal microbiota, which have identified *Firmicutes* (*Bacillota*), *Bacteroidetes* (*Bacteroidota*), and *Proteobacteria* (*Pseudomonadota*) as major components of the bacterial community (25–28). The predominance of these bacterial phyla may reflect their adaptation to the cattle gastrointestinal environment, where microbial communities play a vital role in the fermentation and utilization of complex dietary substrates (29). Operation size explained only 6.5% of the variation in fecal microbial community composition (R² = 0.065, *P* = 0.609), with no statistically significant differences observed among operation-size categories. Similarly, no significant differences in alpha diversity were observed across operation sizes. Variations in diet, feeding practices, management, environmental exposure, or other operation-level characteristics may contribute to microbial community composition independently of operation size. Previous studies have demonstrated that cattle fecal microbial communities can vary with feeding and management conditions. Shanks et al. (25) examined 30 adult beef cattle from six feeding operations (five animals per operation) and reported differences in fecal microbial community composition related to feeding management and fecal starch concentrations. Similarly, Zhang et al. (26) compared six grazing and six feedlot Angus beef cattle and observed differences in fecal microbial community composition between the two feeding systems. Although neither study specifically evaluated cow–calf operation size, their findings suggest that feeding practices and management conditions may influence fecal microbial community composition independently of operation size. The animal group showed a stronger association with fecal microbial community composition than operation size, explaining 20.0% of the variation compared with 6.5% for operation size. Unlike operation size, the difference between cows and calves was statistically significant (R² = 0.200, *P* < 0.001). Cows also exhibited greater observed richness, Shannon diversity, and Simpson diversity than calves. These findings are consistent with the progressive development and maturation of the gastrointestinal microbiota during early life. Hennessy et al. (30) reported that fecal bacterial richness and Shannon diversity increased with calf age, with pronounced changes occurring during weaning and the transition from milk to solid feed. Similarly, Uddin et al. (27) showed that fecal microbial diversity increased from birth to weaning in beef calves and gradually approached the diversity observed in adult cows, whose fecal microbial diversity remained comparatively stable over time. Haley et al. (31) also reported marked differences in fecal microbial community structure between preweaned dairy calves and adult cows. Together, these findings support the greater microbial richness and diversity observed in cows than calves in the present study and suggest that age-related maturation of the gastrointestinal microbiota may contribute to the observed differences in microbial community composition.

Taxonomic differences between cows and calves further supported this developmental pattern. *Lactobacillus* and *Escherichia* exhibited higher relative abundance in calves, whereas *Romboutsia* and *Bacillus* were more abundant in cows. Uddin et al. (27) similarly reported that *Lactobacillus* was prominent in the fecal microbiota of young beef calves during the milk-feeding period and declined as calves aged and forage intake increased, whereas *Romboutsia* became more abundant later in development. These shifts are consistent with the progressive transition from a milk-associated microbial community toward a more mature gastrointestinal microbiota associated with increasing solid feed and fiber intake. The higher relative abundance of *Escherichia* observed in calves may similarly reflect age-related microbial succession, as *Escherichia* is high during early gastrointestinal colonization in calves, especially during the milk-feeding period and declines as calves approach weaning (32, 33). Overall, these taxonomic patterns, together with the greater microbial richness and diversity observed in cows, suggest that animal developmental stages may be an important factor contributing to differences in fecal microbial community composition between cows and calves.

### 4.2 Resistome

The resistome analysis identified a total of 75 unique ARGs belonging to 14 AMR classes in the shotgun metagenomic datasets. Tetracycline resistance genes showed the broadest distribution across operation sizes and animal age groups, followed by aminoglycoside, β-lactam, macrolide, and lincosamide resistance genes. Tetracyclines have historically been widely used in cattle production and remain important therapeutic agents for treating and controlling bacterial diseases, including bovine respiratory and enteric diseases (17). Such antimicrobial exposure can exert selective pressure favoring the maintenance of resistance determinants within cattle-associated microbial communities. However, antimicrobial use alone may not explain the widespread occurrence of tetracycline resistance genes, as these resistance determinants have also been detected in cattle with reduced or no antimicrobial exposure (34, 35). The broad distribution of tetracycline resistance genes observed in the present study is consistent with previous metagenomic investigations of cattle production systems. Noyes et al. (11) reported that tetracycline resistance accounted for a major proportion of resistance-associated sequences detected in cattle production samples, while Zaheer et al. (36) identified tetracycline resistance as a prominent component of the beef cattle fecal resistome. Collectively, these findings suggest that widespread occurrence of tetracycline resistance genes in cattle may reflect both historical antimicrobial selection and the persistence of these genes within the bovine gastrointestinal microbial communities.

Operation size was not significantly associated with overall normalized ARG hit rate, suggesting that operation size alone may not explain the variation in the resistome among beef cow–calf operations. In contrast, calves exhibited a significantly higher normalized ARG hit rate than cows. This finding is consistent with Haley et al. (31) who reported substantially greater ARG abundance in preweaned calves than in adult cows. The greater ARG burden observed in young calves is likely multifactorial. Although antimicrobial exposure can select for resistant bacterial populations, elevated ARG abundance has also been reported in calves without antimicrobial treatment, indicating that antimicrobial use alone may not explain this age-related pattern. Liu et al. (37) associated the elevated ARG burden during early life with the developing gut microbiota, including ARG-associated taxa such as *Escherichia*, and identified maternal colostrum as a potential source of ARGs in neonatal dairy calves. The ARGs detected in colostrum may originate from maternal-associated bacteria or environmental contamination, including teat surfaces and maternal feces, although their precise origin remains uncertain. Together with the differences in microbial community composition and diversity observed between cows and calves in the present study, these findings suggest that age-associated differences in gut microbial ecology may help explain the higher normalized ARG hit rate observed in calves.

ARGs exhibited both shared and unique distributions across operation sizes and animal age groups, indicating that some resistance genes are broadly distributed within the bovine fecal resistome, whereas others showed limited occurrence. Similar heterogeneity in the occurrence and abundance of individual ARGs has been reported across cattle production systems (11, 34, 35). Such variation may reflect differences in microbial community composition, antimicrobial exposure history, diet, management practices, and environmental reservoirs. The greater number of uniquely detected ARGs in calves may also be consistent with the dynamic nature of the developing gastrointestinal microbiome and resistome discussed above. Overall, these findings suggest that the distribution of individual resistance determinants may vary according to host-and operation-level ecological conditions, even when differences in the overall normalized ARG hit rate are not statistically significant.

### 4.3 Mobilome

MGEs play an important role in the dissemination of AMR by facilitating the movement of ARGs within and between bacterial genomes (8). In the present study, 23.9% of the curated ARG hits were located on contigs predicted to be plasmids, indicating that plasmid-associated genomic contexts represented a substantial component of the bovine fecal resistome. Similar proportions were observed across operation sizes and between cows and calves, further suggesting that plasmid-associated resistance determinants were broadly distributed among the cattle groups examined. This finding is consistent with previous studies that have identified plasmid-associated ARGs in the bovine rumen and fecal microbiome (28, 38). Plasmids can carry ARGs and, when transferable, facilitate their dissemination among bacteria through horizontal gene transfer (8). The identification of ARGs on predicted plasmid contigs in the present study suggests a potential for plasmid-mediated ARG dissemination within cattle-associated microbial communities, although actual gene transfer was not demonstrated.

In contrast, ARG hits located on predicted viral contigs were uncommon, accounting for only 0.9% of all ARG hits. This finding is consistent with previous evidence confirming that ARGs are rarely encoded in phage genomes (39), suggesting that predicted viral genomic contexts constitute only a minor component of the fecal resistome identified in the present study. Insertion sequences represented another potential genomic context for ARG mobility, with 7.1% of ARG hits located on IS-containing contigs; however, direct ARG–IS overlap was uncommon (0.95%). IS elements can contribute to the mobilization and rearrangement of resistance determinants through transposition and can also influence the expression of neighboring genes (8, 40). In the present study, most ARG hits located on IS-containing contigs did not directly overlap a predicted IS, indicating that these findings primarily reflect genomic co-localization or proximity rather than evidence of active ARG mobilization. Similarly, although integron-related structures were detected, none of the curated ARG hits were located on integron-associated contigs. Overall, these findings indicate that predicted plasmid-associated genomic contexts were the primary mobility-related feature of ARGs in the fecal resistome identified in this study.

Although this study provides insights into the microbiome, resistome, and mobilome of beef cattle in cow–calf operations, it has several limitations. First, the inclusion of nine cow–calf operations from a single geographic region may limit the generalizability of the findings. In addition, the cross-sectional study design precluded the evaluation of temporal changes in the fecal microbiome, resistome, and mobilome. Detailed antimicrobial-use histories were unavailable, limiting the direct evaluation of associations between antimicrobial exposure and ARG occurrence patterns. Furthermore, short-read metagenomic sequencing limits reconstruction of the genomic context of ARGs; therefore, predicted associations between ARGs and MGEs should not be interpreted as evidence of horizontal gene transfer. Future studies incorporating larger and geographically diverse populations, longitudinal sampling, detailed antimicrobial-use data, and long-read or hybrid sequencing approaches would provide greater resolution of resistome dynamics, genomic context, and the mobility potential of ARGs.

In conclusion, this study demonstrated that animal age group was significantly associated with variation in the fecal microbiome of beef cattle from cow–calf operations, whereas operation size was not. Calves exhibited significantly higher ARG hit rates than cows, whereas no significant differences were observed among operation sizes. Tetracycline resistance genes showed the broadest distribution across operation sizes and animal groups. In addition, a substantial proportion of ARG hits were located on predicted plasmid contigs, highlighting plasmids as an important potential genomic context for ARG persistence and mobility. Overall, this study contributes to the limited knowledge of the fecal microbiome, resistome, and mobilome in beef cow–calf production systems. The observed differences between cows and calves may help ranchers and veterinarians recognize the importance of considering animal groups separately when monitoring antimicrobial resistance (AMR) and reviewing herd health management practices. These findings may also inform targeted surveillance and antimicrobial stewardship efforts in cow–calf operations. Future longitudinal studies involving a larger and more representative number of operations are needed to examine how microbial communities, antimicrobial resistance genes (ARGs), and mobile genetic elements (MGEs) change over time and to investigate the influence of antimicrobial use, animal age, and management practices on AMR distribution and potential transmission.

## ACKNOWELDGEMENTS

The primary author (A.A.) is funded by the “US-Pakistan Knowledge Corridor Scholarship” from the Higher Education Commission of Pakistan and the University of Florida-College of Veterinary Medicine, USA. This work was also supported by funding from the American Rescue Plan (ARP) for One Health Research through the Florida Department of Agriculture and Consumer Services (FDACS), Award No. 16262.

AA: Conceptualization, Methodology, Investigation, Formal analysis, Data curation, Software, Visualization, Writing – original draft, review & editing. LKE: Investigation, Methodology, Writing – review & editing. JHJB: Investigation, Resources, Writing – review & editing. TD: Methodology, Writing – review & editing. ACYL: Writing – review & editing. KCJ: Writing – review & editing. JAH: Methodology, Writing – review & editing. SK: Conceptualization, Methodology, Supervision, Project administration, Resources, Funding acquisition, Validation, Writing – review & editing. All authors read and approved the final manuscript.

## SUPPLEMENTARY MATERIALS

The following supplementary materials are available as additional files:

Supplementary Table S1. Number of sequencing reads generated for 54 pooled fecal metagenomic samples from beef cows and calves across nine cow–calf operations in Florida.

Supplementary Table S2. Phylum-level relative abundance (%) of microbial taxa identified in 54 pooled fecal metagenomic samples.

Supplementary Table S3. Genus-level relative abundance (%) of microbial taxa identified in 54 pooled fecal metagenomic samples from beef cows and calves across nine cow–calf operations in Florida.

Supplementary Table S4. Antimicrobial resistance genes identified in 54 pooled fecal metagenomic samples and their predicted associations with plasmids, viruses, integrons, and insertion sequences.

## ETHICS APPROVAL

Fecal sample collection from beef cattle operations was performed with approval from the University of Florida Institutional Animal Care and Use Committee (IACUC202300000165, approved on 17 March 2023) and in accordance with institutional guidelines.

## DATA AVAILABILITY

The raw shotgun metagenomic sequencing data generated in this study has been deposited in the National Center for Biotechnology Information (NCBI) Sequence Read Archive (SRA) under BioProject accession number **PRJNA1544148.** Additional data supporting the findings of this study are provided in the article and its Supplementary Materials.

